# Diverse genetic origins of medieval steppe nomad conquerors

**DOI:** 10.1101/2019.12.15.876912

**Authors:** Alexander S. Mikheyev, Lijun Qiu, Alexei Zarubin, Nikita Moshkov, Yuri Orlov, Duane R. Chartier, Igor V. Kornienko, Tatyana G. Faleeva, Vladimir Klyuchnikov, Elena F. Batieva, Tatiana V. Tatarinova

**Author notes:** Corresponding authors: AM, TT. These authors contributed equally to this work (Alexander S. Mikheyev, Tatiana V. Tatarinova).

## Abstract

Over millennia, steppe nomadic tribes raided and sometimes overran settled Eurasian civilizations. Most polities formed by steppe nomads were ephemeral, making it difficult to ascertain their genetic roots or what present-day populations, if any, have descended from them. Exceptionally, the Khazar Khaganate controlled the trade artery between the Black and Caspian Seas in VIII-IX centuries, acting as one of the major conduits between East and West. However, the genetic identity of the ruling elite within the polyglot and polyethnic Khaganate has been a much-debated mystery; a controversial hypothesis posits that post-conversion to Judaism the Khazars gave rise to modern Ashkenazim. We analyzed whole-genome sequences of eight men and one woman buried within the distinctive kurgans of the Khazar upper (warrior) class. After comparing them with reference panels of present-day Eurasian and Iron Age populations, we found that the Khazar political organization relied on a polyethnic elite. It was predominantly descended from Central Asian tribes but incorporated genetic admixture from populations conquered by Khazars. Thus, the Khazar ruling class was likely relatively small and able to maintain a genetic identity distinct from their subjugated populations over the course of centuries. Yet, men of mixed ancestry could also rise into the warrior class, possibly providing troop numbers necessary to maintain control of their large territory. However, when the Khaganate collapsed it left few persistent genetic traces in Europe. Our data confirm the Turkic roots of the Khazars, but also highlight their ethnic diversity and some integration of conquered populations.

## Introduction

Steppe nomad conquests over the past two thousand years repeatedly upended the history of settled Eurasian peoples. These included wars fought between the Xiongnu and the Chinese at the dawn of the common era, and the European invasion by the Huns in the early Middle Ages. Afterward, numerous other tribes, most notably the Mongols, fought their way out of Central Asia and set up relatively short-lived Khaganates in the occupied lands. Unfortunately, except for short runic inscriptions, these nomads left few contemporary written records. Much of what we know about them comes from accounts composed by their traditional enemies. Contemporary animosity skews existing accounts, and descriptions of many historical events, such as the number of soldiers killed in battle. Exaggerations were introduced either for dramatic effect or for the purposes of contemporary propaganda. Furthermore, steppe nomad societies typically consisted of many allied tribes and ethnicities, and contemporary outsiders struggled to understand their function and organization. As a result, little is known about the ethnic composition of steppe nomads or their relationships with present-day populations or how they may have interacted with local populations during the period of occupation. No single population of steppe nomads yet been studied in detail to understand its ethnic composition and relationship with present-day populations.

The Khazar Khaganate was a particularly important polity, which lasted from the VII to the IX centuries AD and laid between the Caspian and Black Seas (Figure 1A). It formed a buffer between the Byzantine Empire, the Umayyad Caliphate, and other steppe nomads. Its longevity produced a more detailed archaeological record than other Khaganates and provides a window into the structure of steppe nomad societies. Although the Khazars dominated trans-Eurasian trade routes and maintained extensive long-term contacts with settled societies in Europe and the Middle East, their origins and ethnic composition remain controversial. No indigenous records of their language exist, and though many of the Khazar words appear to be Turkic in origin, few of them have reached the present day, making detailed paleo-linguistic analysis difficult. Also, no present-day populations with Khazar descent documented by historical records exist. Overall, we know that the Khazar Khaganate was polyethnic, multilingual and professed all major monotheistic religions as well as Tengrism, but little about the actual ethnicities constituted it, particularly the ruling classes.

**Figure 1:**
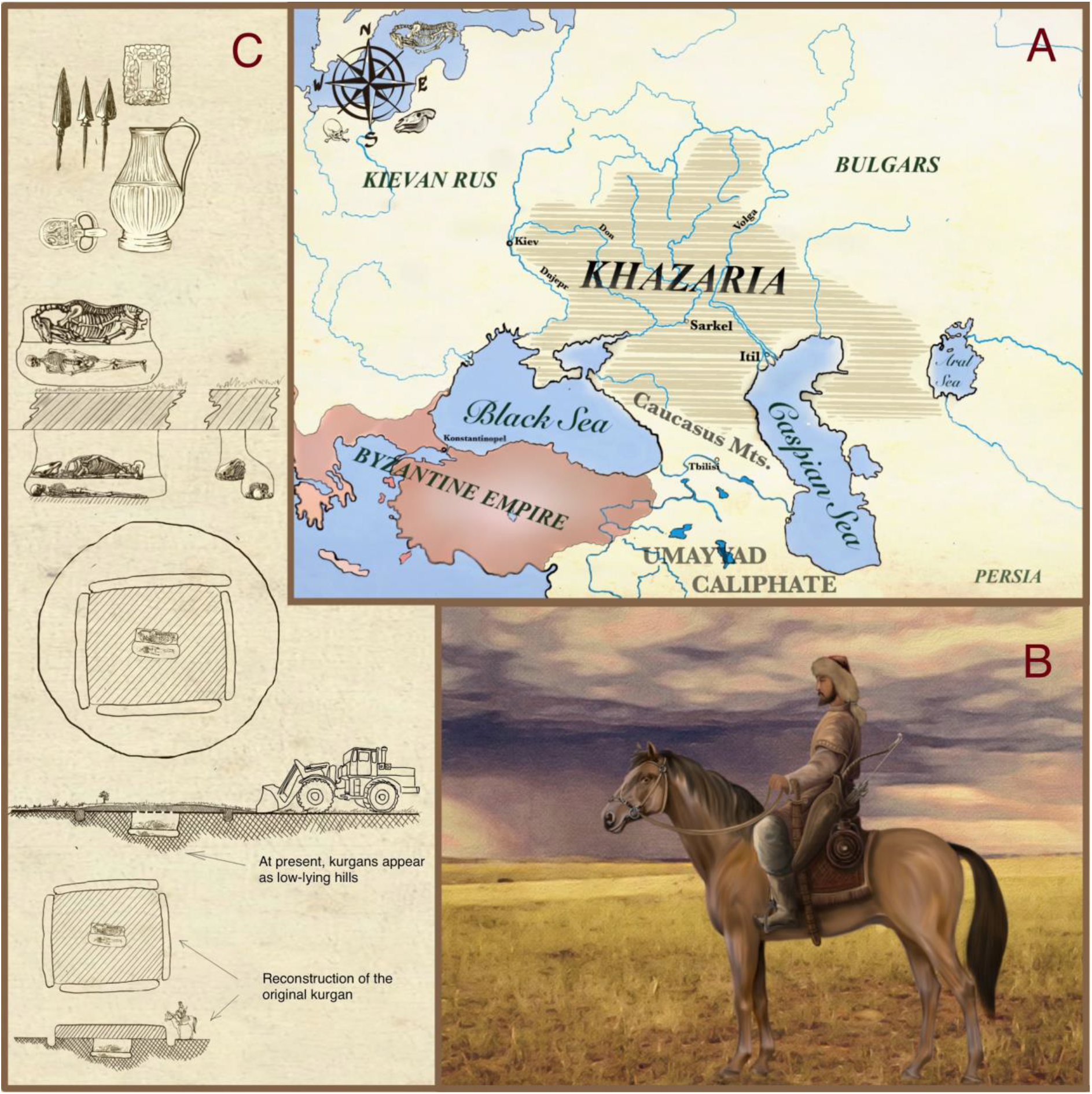
The Khazar Khaganate and its archaeological history. (A). The Khaganate adjoined the Turkic tribes to the East, the Umayyad Caliphate to the South and Byzantine Empire to the West, warring, and trading with all of them throughout its history. The boundaries of the Khaganate are approximate and shifted over the course of its existence. (B). Reconstruction of a Khazar mounted archer, with mount and traditional weaponry including the recurved compound bow and sword, and equestrian gear including bridle and stirrups, which were typically included in Kurgan burial sites (C). Top: As a result of centuries of soil erosion and plowing, the present-day kurgans appear as low-lying hills, with one or more graves inside them. Characteristic square trenches often used for ritual animal sacrifices surrounded Khazar burial sites. Typically, the horse is buried alongside the warrior on a separate ledge. Common grave goods included weapons, ornaments, coins, and pottery. Bottom: Artistic rendering of the kurgan during Khazar times.

The reliability of contemporary accounts made by travelers is likewise in doubt, as they are filled with contradictions, which may have been introduced by subsequent transcriptions and translations. For example, the Persian geographer al-Istakhri who traveled in the Khazar lands described two different ethnicities of Khazars: fair-skinned Ak-Khazars and darker-skinned Kara-Khazars (excerpts from al-Istakhri’s works [1,2]). However, according to some translations of his work he also noted that Khazars did not resemble Turkic peoples [1]. Adding to the confusion, al-Istakhri used Turkic words for “white” and “black” (i.e., “ak” and “kara”) to name the two Khazar ethnicities. Khazarian toponyms have words that can be classified as Turkic, as well as not Turkic [3]. Historical records do surprisingly little to clarify the Khazars’ ethnicity, but rather add to their mystery [1].

The fact that some Khazars were described as fair-skinned by contemporaries, together with the possible adoption of Judaism by the state and by the ruling classes in the IX century, raised the hypothesis that conversions among the Khazars gave rise to Ashkenazi Jews (the ‘Khazarian hypothesis’). However, archaeological data do not support the adoption of Judaism by a large fraction of the Khaganate, as very few items with Judaic symbology have been found [4, 5]. It seems most likely that only some representatives, if any, of the ruling elite underwent the religious conversion, while the exact timing of the purported Judaism adoption also remains a debatable issue [6]. Nonetheless, under the Khazarian hypothesis, rather than being dispersed by the southward advance of the Kievan Rus’ tribes, the Judaized Khazars migrated to Europe and formed the genetic core of modern-day Ashkenazim. This hypothesis has been hotly debated for two centuries, initially using historical material, but also increasingly, using genetic studies. The existence of conflicting reports has led to debates about their veracity and authenticity; even the very practice of Judaism by the Khazars has been questioned [7]. Yet, recent work on geographical genetics and linguistics of present-day Ashkenazi populations lent support to the hypothesis of the Khazarian origin [8–10]. However, the methodology of this work has been subjected to critique, citing the difficulty of reconstructing ancestral Jewish populations without appropriate Khazarian proxies in using present-day data. The genetic homogeneity of the worldwide Jewish population is also problematic [11, 12]. Consequently, given the paucity of historical records, and the complexity of migration patterns by Jews and the nomadic steppe tribes, this hypothesis is only testable by examining evidence of genetic relatedness between Ashkenazim and the Khazars using genomic analysis of archaeological remains.

The Khazar Khaganate was a polity that united numerous peoples with different ways of life [13]. It appears that ethnic Khazars comprised a relatively small proportion of the ethnically diverse society, which was made up of settled agriculturalists, town workers, merchants, as well as steppe nomadic herders. One of the uniting archaeological criteria that characterized Khazaria is the Saltovo-Mayatskaya culture [14], which was a shared assortment of items and tools used by different peoples inhabiting the Khaganate. Among those items, the most important and numerous are the ceramic vessels in the Alan or North Caucasus style, that were in common use by both nomads and farmers. Consequently, the widespread popularity of these items among diverse ethnic groups makes it difficult to associate them with any single group. This difficulty extends to the identification of the Khazars themselves using the artifacts alone.

In the past the Khazar graves were difficult to identify among the numerous burial sites, settlements and cities excavated by archeologists [15]. However, from the middle of the 1970s, excavations along the Lower Don river uncovered numerous kurgans with square trenches, which were almost unknown beforehand [16, 17]. Many of these kurgans were destroyed by looting or plowing, and how they looked initially is difficult to completely reconstruct, but their large number and common characteristics allowed a reliable overall picture to emerge.

Inside each kurgan, there was typically one burial with square trenches that surround the burial mound. Male kurgans are much more common than female kurgans [18], and they frequently include remains of weapons, such as arrowheads and bone plates that made up part of the warrior’s composite bow. Buried in the kurgan near that warrior there are usually bones of a horse or sometimes an entire horse skeleton complete with bridle, stirrups and other tacks. While the majority of the kurgans have been looted, even what remains attests to the relative wealth of those buried there: burial goods include golden Byzantine coins, silver belt buckles, decorations, and even expensive imported silver vessels (the richest burials were described in several publications [19, 20]). Notably, not a single item associated with the practice Judaism was ever recovered from these burial sites.

The common archaeological features of these burial sites, including the weaponry and wealth, indicated that the deceased belonged to a single ethnic and social group within the Khaganate. They were clearly the military and political elite of the Khaganate, which were the Khazars themselves. As a result, there is a consensus among historians that kurgans with square trenches are specifically Khazar burial grounds [21]. At the same time, the ethnicity of the Khazars remains hotly debated. Many lines of evidence suggest their Turkic origins [22], including analysis of their runic inscriptions, and the hypothesis that they were Turkic refugees fleeing the collapse of the Western Turkic Khaganate [15], which was destroyed in the middle of the VII century due to internal strife, as well as the onslaught of the Tang Empire. An interesting point of view on the events of this period is given by Flerova, who linked the Khazar burial practices to those found in Central Asia, and noted that they appeared along the Don river immediately after the fall of the Western Turkic Khaganate [23]. To conclusively resolve centuries of debate, we believe that the analysis of genetic data can be decisive in resolving the genetic affinity of the Khazars.

Here we explore the ethnic composition of the Khazars, and also specifically test the hypothesis of their relatedness to contemporary Ashkenazi Jews, by sequencing nine genomes from Khazar kurgans in southern Russia. Based on physical anthropological investigations, these burials belong to a range of ethnic types and provide a window into the genetic makeup of the society. We found that the Khazar elite drew from a variety of Central Asian tribes with a mostly Turkic genetic composition and with some Siberian and East Asian components, as well as notable Caucasian and Middle Eastern genetic contributions to at least some of their members. We found no significant trace of Ashkenazi genetic composition in either nuclear, mitochondrial or Y-chromosome data, strongly indicating that Khazars were generally not related to them. These findings support the view that the Khazars were a mixture of predominantly Turkic steppe tribes, who conquered westward territories in probably the same manner as Scythians, Huns, Pechenegs, and Mongols before and after them.

## Results

Assignment of individuals to populations, particularly with genetic admixture, and particularly using low-coverage data is challenging, therefore we used two independent methodologies to confirm our results based on nuclear data analysis. Our main analysis was based on a method that explicitly accounts for genotype uncertainty using likelihoods derived from potentially low-coverage data and allows bootstrapping of data to confirm admixture estimates (see *Analysis using genotype likelihoods)* [24]. We also used a combination of well-established Admixture, GPS, and reAdmix methods using a separate database to examine the geographic origins of individuals while accounting for admixture (see *Analysis using admixture vectors)* [25, 26]. This analysis was conducted using present-day data and data from Bronze-age populations from the first century BC to control for any geographical changes that may have occurred in the past three thousand years. We found that these different approaches produced a consistent assessment of Khazar provenance.

### Sequenced data reference populations

Previous work has shown that most populations in the region of interest, which includes much of Central Asia, the Middle East, and Eastern Europe, are structured geographically [27]. While there is evidence of admixture from Turkic-speaking populations into non-Turkic ones in the 9th—17th centuries, most of the population structure is much older [27]. Therefore, data from the present-day population should be geographically informative for Khazars. Therefore, to understand Khazar origins, we synthesized a reference panel of 1,933 individuals representing 56 Eurasian populations from previously published data that were genotyped using Illumina Infinium arrays [11, 28], which intersected at 120,278 polymorphic sites.

We generated whole-genome data on nine individuals, eight men and one woman (Table S1). These data were generated at low depth (Table S1) producing anywhere from 2,400 to 39,315 covered sites intersecting with the Geno 2.0 reference database [25]. This database represents 98 worldwide populations and subpopulations with 15 samples per population on average (corresponding to 615 unrelated individuals), genotyped on Illumina’s GenoChip array. This array contains 130,000 autosomal ancestry informative markers. Admixture analysis [29] has previously identified nine admixture components corresponding to nine worldwide source populations [25]. It was supplemented with the Eurasian dataset [28], adding 1076 samples collected in different parts of Russia, Kazakhstan, Georgia, Uzbekistan, and Kyrgyzstan. These samples were genotyped on the Illumina Infinium 370-Duo, 370-Quad, or 610-Quad arrays. In order to add these samples to the reference database, we have extracted 130,000 Geno 2.0 ancestry informative markers and applied Admixture [29] analysis in supervised mode was conducted to convert SNPs to the 9-dimensional admixture vectors [25].

### Genetic relationships with present-day populations

#### Admixture analysis using genotype likelihoods

We also found that despite the history of steppe nomadic migration and conquest, their genetic relationships closely reflect present-day geographic location (Figure 2A). Thus, these populations are strongly informative about the geographic origins of the Khazars. By projecting the nine Khazars onto the reference panel, we find that three of them had predominantly Asian genetic ancestry, without significant European contributions, while the other two had significant percentages of genes from the Caucasus region, in addition to those from Eastern Russia and Asia (Figure 2B). The mitochondrial haplotypes matched nuclear findings, with Asian haplotypes in the three Khazars that had an affinity to present-day Asian ethnicities, and European haplotypes in the others (Figure 3C). Finally, these results matched previous physical anthropology findings, which classified the Khazars into Caucasoid or Mongoloid racial types [30] (Figure 3D). These results confirm that the Khazar elite was polyethnic, composed of Asian steppe nomads, and individuals with recent European ancestry admixed with Asian nuclear genetic components.

**Figure 2:**
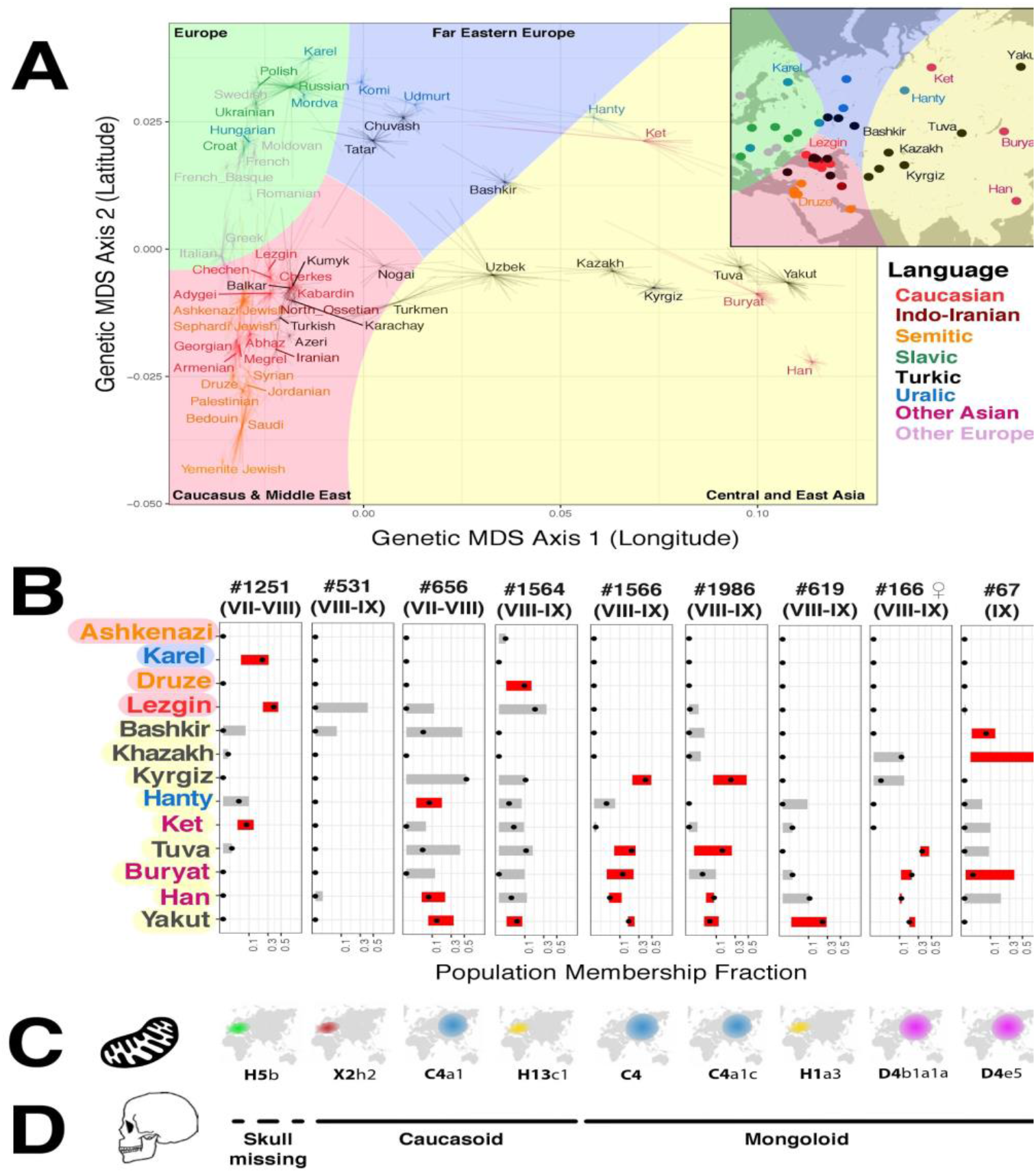
56 present-day populations and their relationship to the Khazar elite. (A) Genetic distances between modern Eurasian populations correspond to their geographic origins. The main plot shows the first two multidimensional scaling components based on genetic data, which roughly correspond to longitude and latitude in geographic space (inset). Each individual sample is represented by a line connected to a population centroid, and populations are color-coded by language category. Despite the extensive history of migration in Central Asia, the correspondence genetic data and between present-day geographic location suggest that genes can act as a proxy for geographic ancestry. (B) Projection of the nine Khazar specimens onto the present-day population suggests polyethnic origins from Siberia and East Asia and the Caucasus. The admixture proportion for each population is shown as a range based on bootstrapping the data, with proportions significantly different from zero in red. Three individuals (#1251, #1564, and #531) show a mixture of Asiatic and Caucasian components, while the other six show largely Asian genetic components. Notably, significant traces of Ashkenazi Jewish admixture are absent from each of the individuals. Each specimen is labeled by century, and the female specimen #166 is indicated by 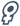. (C) Mitogenomes correspond to the nuclear origins of the Khazars. The individuals with more European genomic DNA have mitogenomes typically found in the Caucasus and elsewhere throughout Europe, while the others have typically Asian haplotypes. (D) Previously published physical anthropology data match genetic data in the four cases where skulls were available. We predict that #1251, where no skull was found was Caucasoid in appearance. Though many Khazars descended from Turkic tribes, some originated from other Siberian or East Asian peoples, and some had admixture from the Caucasus or the Levant. Nonetheless, their genetic makeup remained distinct from that of the conquered populations.

#### Analysis using admixture vectors

The GPS algorithm determines a single point on the map that corresponds to the location of individuals with the most similar genotype (measured as a Euclidean distance in the 9-dimensional space of the “ancestral” admixture components). reAdmix algorithm [26] presents an individual as a weighted linear combination of reference populations. Both tools use admixture vector representation of genotype data. We have extracted ancestry informative markers [25] from the nine Khazar samples (using positions with Q>=20 and depth of coverage >=2) (see Table S4 for the number of sites). For the purpose of comparison with previous studies, we retained the names of the components, although they may be an accurate representation of ancestry. ADMIXTURE analysis (Figure SF3 and Table S5) shows that the nine samples differ in amounts of North East Asian, Northern European and Mediterranean components: six of them have a significant North-East Asian component, while the other three have more South West Asian, Mediterranean, and Northern European ancestry. This agrees with the prevailing assumption of the genetic diversity of Khazarian society.

### Khazars in the context of Bronze-age material

From the published literature (see Materials and Methods) we added 587 ancient Eurasians (1 BCE 0 3000 BCE) o the GPS/reAdmix database used above to understand possible relationships between the Khazars and Bronze Age populations. We found that most of the Khazar samples grouped together (Tables S6-S8, Figure SF4), and that “Pazyryk Iron Age” samples consistently appear as reAdmix matches for the Khazar samples. Pazyryk is a Scythian period nomadic culture (6th–3rd centuries BC [31]) that flourished in Southern Siberia, and several well-preserved burials containing mummified remains were found in Altai mountains. However, with the genetic resolution of our data, both present-day and Bronze-age material suggests that the geographic origins of the Khazar populations were geographically similar (Figure 3). In other words, both data sets point to the Khazar origins across a broad swath of Central Asia and admixture with Eastern Europe and the Middle East.

**Figure 3:**
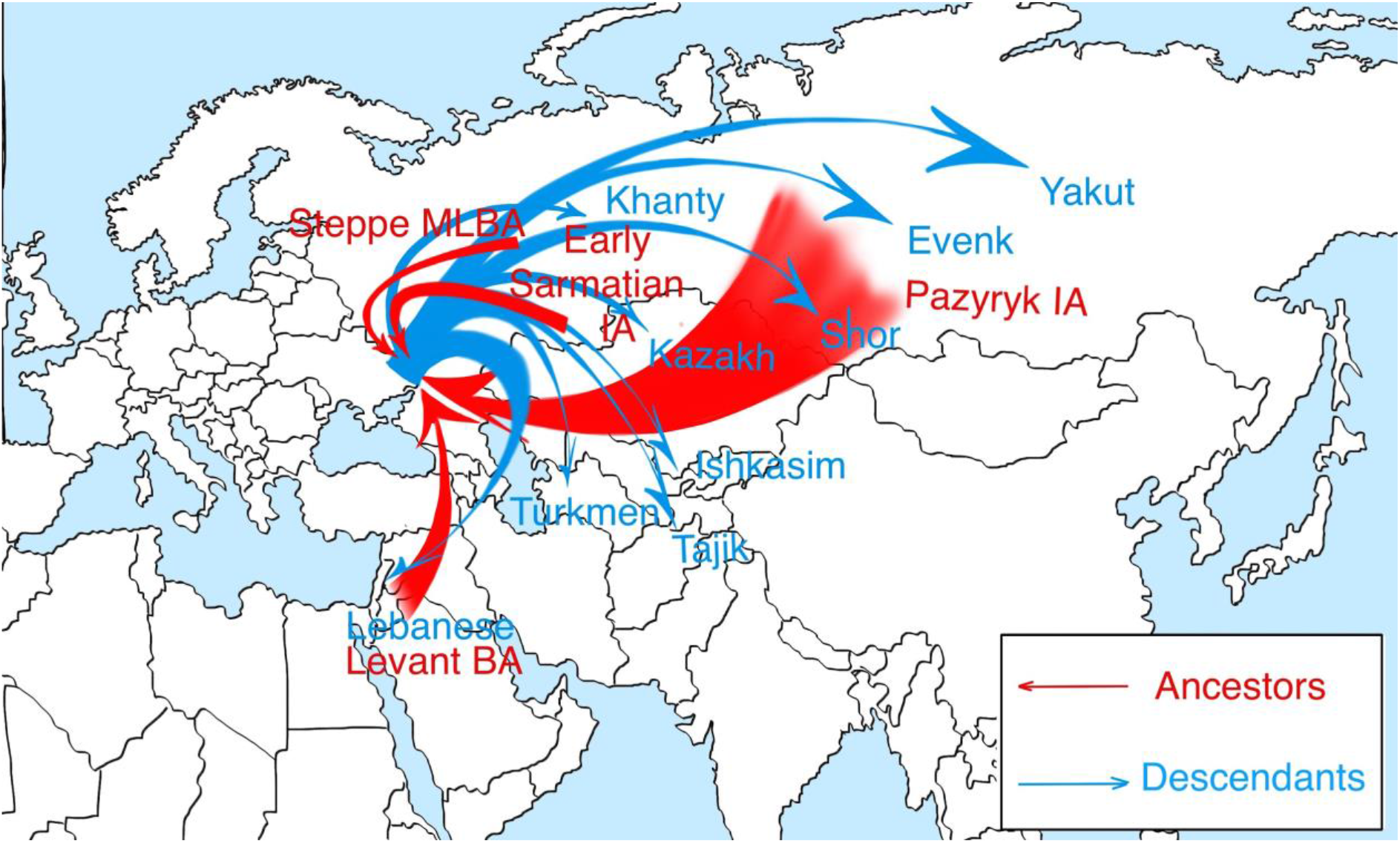
Visualization of reAdmix results using Bronze Age and modern samples (“Ancestors” and “Descendants”). Red arrows point from the inferred geographic locations of Bronze Age populations represented in the Khazar elite. Blue arrows point towards present-day populations. Names of closely related populations are highlighted on the map. The width of lines represents the strength of the inferred relationships. Overall, present-day and Bronze age data indicate that Khazars were of largely Central Asian origin, with admixture from either the Caucasus or the Near East. In other words, the Khazars were largely similar to populations located in East Asia both in the Bronze age and in the present day.

## Discussion

Our analysis sheds light on the centuries-old mystery of Khazar origins providing an insight into the structure of steppe-nomad confederations and how they interacted with conquered populations. Our data support historical accounts of other steppe nomad groups, such as the Mongols, and previous archaeological and historical inference about the Khazars. The Khazar and the Mongol governments were both confederations of different steppe tribes, drawn over an impressive geographical range and a wide variety of ethnicities. While Turkic tribes likely formed their core, it is probable that they drew members from present-day Siberian and East Asian populations. As a result, they likely spoke a mixture of languages, though it seems logical that they would have relied on a Turkic base, consistent with Turkic roots of the few Khazar words that survived to the present day and the runes associated with their culture. Furthermore, the burial practices of the Khazars are also similar to those of Turkic sites in Central Asia [32].

We can infer several conclusions about the social structure of the Khazars ruling class based on the entirety of our data, archeological findings and contemporary accounts, which are consistent among themselves. While the Khazars were polyethnic, it appears that they nonetheless had a race-based class system, with Asiatic conquerors at the top. The contemporary geographer Al-Istakhri wrote that “light-skinned” Khazars descended from slaves [1, 33, 34].

According to a previous survey of Khazar burials, sixty-five percent of male kurgans were occupied by Mongoloids, and thirty-five by Caucasoids [18]. Both Caucasoid and Mongoloid men were buried in kurgans with full honors (Figure 2), suggesting that they had a privileged status within the Khazar society. If we consider men buried with Caucasoid admixture to be ethnically alien to the Khazars (or born to a mixed-race family), there was some possibility of social mobility for newcomers, regardless of their ethnic origin. It is extremely interesting that the overwhelming majority of women (more than 90%) buried in mounds with trenches are Mongoloids [18]. Perhaps the Mongoloid women, being closer genetically related to the Khazars, belonged to a more privileged part of society and were more desirable as brides. This is further supported by the observation that none of the males in our sample had purely European or Middle Eastern ancestry. Even the four morphologically or genetically Caucasoid Khazars (samples #656, #533, #1251, and #1564) had some mixture of Asiatic and European nuclear DNA. This suggests that Khazar or other Asiatic ancestry may have been a major social asset in this society.

The origins of the admixed individuals are even more speculative. There are extensive accounts of steppe nomads marrying or abducting local women [35], and this was likely the practice of the Khazars as well. It is possible that mixed-blood descendants could have achieved a relatively high rank in society, at the very least meriting burial with full military honors, but whether they could marry Khazar women, and become fully integrated into society remains unknown. Writing about the Mongols several centuries later, Giovanni de Carpini wrote that children as young as two or three were trained in horseback riding and archery, making it likely that adopting the steppe nomads warrior skill set required being born into that culture [36]. However, in our Khazar sample, there is one individual with Caucasoid facial features and an Asiatic mitochondrial haplotype (#656). He could have been a child of a mixed marriage between a woman of Central Asian descent and a non-Khazar (or a mixed-blood Khazar). Future genetic investigation of burial sites not associated with the elite should shed light on the relationship between the Khazars and their thralls, as well as opportunities for upward mobility in the Khaganate. Nonetheless, it appears that the Khazar elite was composed of Asiatic steppe nomads which remained remarkably genetically and culturally distinct from the peoples they conquered. One possible exception to this observation might be the Bulgars, who were absorbed into the Khaganate when Old Great Bulgaria was conquered by the Khazars in VII century AD and may have been assimilated. However, given the common Turkic genetic background of the Bulgars and Khazars, these ethnicities may be difficult to tell apart either archaeologically or genetically.

Genetic data allow us to conclusively answer the question of whether Judaized Khazars may have migrated to Eastern Europe to give rise to the present-day Ashkenazim, as proposed by the Khazarian Hypothesis. This scenario seems highly doubtful in the face of genomic data based on four lines of evidence. First, no significant Ashkenazi genetic affinity was detected in any of the sequenced individuals (Figure 2B). Second, all of the studied Khazars, even those with significant Caucasian ancestry, had significant Asiatic nuclear genetic contributions, which are missing from present-day Jewish populations [11]. Third, while local women were recruited into Ashkenazi communities, none of the identified mitochondrial haplotypes are present in present-day Ashkenazi Jews [37, 38]. Finally, the European genetic components of the Khazars derive from the Caucasus tribes that were under control of the Khaganate, rather than from more distant Levantine populations more closely related to Ashkenazi and Sephardic Jews (Figure 2). While Jews lived in the territory of the Khazar Khaganate along with Christians, Muslims, and pagans [1], it seems unlikely that they formed its ruling classes, which were dominated by steppe nomads from the East, and thus the Khazars could not have been the progenitors of the Ashkenazim.

Our investigation indicates that over their centuries of residing in Eastern Europe, the Khazars retained many Central Asian characteristics. These include traditions, including burial practices and the traditional nomadic lifestyle, and strong genetic affinities to Turkic peoples among the Khazar ruling class. While mixed-blood Khazars could join its warrior class, our genetic analysis is consistent with previous anthropological investigations that found that Khazars were distinct from endemic inhabitants of Southern Russia and the Caucasus and remained so over the course of several centuries. Our analysis suggests that when the Khazar Khaganate collapsed, it did not go on to form a distinct population that exists in the present day. Instead, its elite was either destroyed, was assimilated by other Eurasian populations, or fled to Central Asia.

## Materials and Methods

### Materials

The nine skeletons were collected from typical Khazar burial kurgans in the southern Russian steppes (see Supplement for details). Genotypes of ancient Eurasians (1st millennium BC-10th millennium BC) were obtained from https://reich.hms.harvard.edu/datasets; these datasets correspond to the following recent publications: [26, 39–50]. The datasets were stratified into 6 cohorts by the estimated age of the samples: 0-1000 BC, 1000-2000 BC, 2000-3000 BC, 3000-4000 BC, 4000-5000 BC, and above 5000 BC. We used worldwide reference data from several sources [11, 25, 28, 51] to compile a dataset of modern populations. The datasets were converted into suitable formats for NGSadmix [26, 52], reAdmix [26], and TreeMix [53] analyses.

### Sample preparation

DNA was extracted in the specifically prepared clean room. Bone fragments were extracted using a rotary autopsy saw. These fragments were left in running water for 15 minutes, then the top plaque was removed with a sterilized scalpel and a brush, and the bone fragments cleaned with a cleaning solution, bleached, and washed in deionized water. Then the fragments were UV treated for 15 minutes on each side and dried in a shaker-incubator. Bone surfaces were cleaned with a portable dental drill using separate sterile tips for each bone. The bone powder obtained from the surface of the sample was discarded. A new sterile drill tip was used to obtain bone powder from inner layers of bones and transferred into a separate sterile 15 ml centrifuge tube for DNA extraction. In order to extract DNA from the powder, we used 1-2 g of bone powder and added 10 ml of sterile 0.5M EDTA solution. The supernatant was discarded, and the cycle was repeated twice. 3.5 ml of lysis buffer, 150 μl of proteinase K and 300μl of 1M DTT solution were added to the remaining sediments. The samples were thoroughly mixed, then incubated at a temperature of +56°C for 2 hours and then at +40°C for 16 hours. Afterward, the samples were centrifuged at 5000g for 10 minutes. The supernatant was transferred to separate sterile 15 ml centrifuge tubes. An equal volume of a mix phenol/chloroform/isoamyl alcohol (25:24:1) was added to the supernatant and vortexed at high speed for 30 seconds. The tubes were then centrifuged for 10 minutes at 5000g. The topwater phase was carefully transferred into sterile 15 ml centrifuge tubes, without touching the interphase. The procedure of phenol extraction was conducted twice more. An equal volume of n-butanol was added to the aqueous solution containing DNA, vortexed vigorously for a couple of seconds, then the tubes were centrifuged at 5000g. Afterward, the top layer of butanol was removed, leaving an aqueous DNA solution in the tube. The final volume of purified DNA samples was approximately 100μl.

### Library preparation and Sequencing

We have followed the library preparation and “U-selection” protocol of Gausauge and Meyer [54, 55] which exploits one of the most distinctive features of ancient DNA—the presence of deoxyuracils—for selective enrichment of endogenous DNA against the background. “U selection” extracts up to five times more of endogenous DNA sequences. The sequencing was conducted on Illumina HiSeq 2500 using single-end sequencing.

### Sequencing reads processing

Fastq reads were analyzed with FastQC [56], and adapters were trimmed using the *cutadapt* tool [57]. The trimmed reads were aligned to the Genome Reference Consortium Human Build 37 (GRCh37) using *bowtie2 [58]* mode *--very_sensitive_local.* MapDamage [59] was used to assess the degree of specific post-mortem modifications, and ANGSD [60] to estimate contamination levels in the analyzed genomes, using the X chromosome that exists in one copy for male samples. Variants were called using the *gatk [61]*, with *samtools [62]*, *vcftools [63]*, and *plink2 [64, 65]* was used for data type conversion and subsetting.

### Data preparation and pre-processing

MDS analysis was performed with *plink[64, 65]*. MT haplogroups were determined using the BAM Analysis Kit (http://www.y-str.org/2014/04/bam-analysis-kit.html).

ADMIXTURE [29, 66] was used to conduct unsupervised and supervised admixture analysis. NgsAdmix [52] was used for finding admixture proportions from NGS data based on genotype likelihoods. 100 bootstrap replicates were conducted for each analysis. GPS [25] was applied for approximate geo-localization of ancient samples, and reAdmix [26] was used to represent Khazar samples as a weighted sum of present-day and contemporary individuals. A reference dataset for GPS and reAdmix was obtained from Elhaik et al [25]. This database represents 98 worldwide populations and subpopulations with 15 samples per population on average (corresponding to 615 unrelated individuals), genotyped on Illumina’s GenoChip array (Geno2.0). Geno2.0 array contains 130,000 autosomal ancestry informative markers. Admixture [29] algorithm in supervised mode was used to convert the Khazar genotypes to the 9-dimensional admixture vectors of “ancestral” proportions (“North-East Asian”, “Mediterranean”, “South African”, “Southwest Asian”, “Native American”, Oceanian”, “Southeast Asian”, “Northern European”, “Sub-Saharan African”). For the purpose of comparison with previous studies, we retained the names of the components, although they may not be an accurate representation of ancestry. The Geno 2.0 database was supplemented with the Eurasian dataset [28], adding 1076 samples collected in different parts of Russia, Kazakhstan, Georgia, Uzbekistan, and Kyrgyzstan. These samples were genotyped on the Illumina Infinium 370-Duo, 370-Quad, or 610-Quad arrays. In order to add these samples to the reference database, we have extracted 130,000 Geno 2.0 ancestry informative markers and applied Admixture [29] analysis in supervised mode was conducted to convert SNPs to the 9-dimensional admixture vectors [25].

### GPS and reAdmix

To infer the geographical coordinates (latitude and longitude) of an individual given the K admixture frequencies, GPS requires a reference population set of N populations with both K admixture frequencies and two geographical coordinates (longitude and latitude). SNPs corresponding to 130K Ancestry Informative Markers [26, 67] were extracted using PLINK [64, 65] and ADMIXTURE tool [29] in supervised mode using the reference populations described in Elhaik et al. [25]. The genotype data were converted into 9-dimensional vectors, and then Geographic Population Structure (GPS) [25] and reAdmix [26] algorithms were used to infer the provenance of the samples and to confirm self-reported ethnic origin. For each tested individual, the GPS algorithm determines a location on a world map, where people with similar genotypes are likely to reside. GPS predictions were followed by analysis with reAdmix, which models an individual as a mix of populations and permits user-guided conditional optimization. The GPS algorithm was used to determine a single point on the map that corresponds to the location of individuals with the most similar genotype (measured as a Euclidean distance in the nine-dimensional space of the “ancestral” admixture components). reAdmix was used to represent the studied individuals as a weighted linear combination of reference populations. Both tools use admixture vector representation of genotype data. It is possible to create admixture vectors using two strategies - supervised and unsupervised. Both modes are implemented in ADMIXTURE [29]. We have extracted ancestry informative markers [25] from the nine Khazar samples (using positions with Q>=20 and depth of coverage >=2) and 587 Ancient Eurasian (1 BCE-3000 BCE) samples from the sources listed above.

### TreeMix analysis

We used the three-population test (f3) developed by Patterson, et al. [68], implemented as the *threepop* program of the TreeMix [53] package. Statistic f3 (O; A, X) measures relative amount of genetic drift shared between the test population A and a reference population X, given an outgroup population O distant from both A and X. Outgroup f3 statistic is always positive, and its values can be interpreted only in the context of the reference dataset X. The statistical significance of f3 was assessed using a Z-score: a statistic divided by its standard deviation. We reported combinations with the absolute value of Z-score above 3; this threshold corresponds to a p-value of 0.00135 and is commonly used for similar applications [28]. The three-population statistics were calculated in the *threepop* software included in Treemix with the block-size parameter *(-K)* set to 100 for all samples. The choice of *K* is determined by the linkage disequilibrium and the density of SNP.

## Funding information

Dr. Kornienko was supported by the state assignment projects (# AAAA-A19-119011490038-5), “Kalmyk Research Center of the Russian Academy of Sciences”, Elista, Russia 358000 and (# 01201363186), Federal Research Center, Southern Scientific Center of the Russian Academy of Sciences, Rostov-on-Don, Russia, 344006.

## Acknowledgment

The authors express gratitude to Mr. Michael Christie for an artistic rendering of Khazars and burials.

## Author Contributions

VK and EB have described the samples, IVK and TGF has extracted DNA and conducted haplogroup analysis, LQ and AM conducted sequencing, ASM, AZ, NM, UP and TVT analyzed data, DRC prepared the draft of the manuscript, TVT and ASM have designed the study, and jointly supervised all aspects of this work. All authors have read and approved the manuscript.

## Competing Interest Declaration

The authors declare no competing interests.

## Data availability

Sequencing data have been submitted to the DNA Data Bank of Japan and will be released upon publication.

## Additional Information

Supplementary Information is available for this paper.

Correspondence and requests for materials should be addressed to ASM and TVT

Reprints and permissions information is available at www.nature.com/reprints

## Extended data figure and table legends

Figure SF1: Location of the kurgans

Figure SF2: Artistic rendering of stepped (A) and niche graves (B).

Figure SF3: Admixture profiles of Khazar samples together with other ancient DNA samples.

Figure SF4: Clustering of ancient samples. Six out of nine Khazar samples cluster together (green cluster)

Table S1: Samples description showing gender, age, date, location, and sequencing statistics (bases covered, average coverage, a fraction of the genome covered by at least one read).

Table S2: Contamination. The table lists the results of ANGSD for 9 Khazarian samples. Two estimators are used: Methods of Moments (MoM) and Maximum Likelihood (ML). Also, the analysis can be conducted discarding (right column) and not discarding (middle column) X chromosomal regions with low mappability 7. Contamination level is computed by 4 methods (Method 1(old/new) and Method 2(old/new), using the code from Rasmussen et al 7).

Table S3: Mitochondrial haplogroups. MT age estimates were taken from Behar et al.10 and Derenko et al. 11.

Table S4: The number of Ancestry Informative Markers based on the sequencing quality cutoffs. The insufficient number of markers retained for DP3 is typical for aDNA studies.

Table S5: Admixture profiles of Khazar samples for K=9

Table S6: Summary of GPS results, indicating the nearest reference population and Euclidean distance between the sample and reference admixture vector. A small distance indicates that the reference population is a good match for the sample vector. Large distance may indicate either the absence of a suitable reference population or mixed origin of the sample.

Table S7: reAdmix using a database of modern individuals. A maximum of four source populations was allowed for each sample. When proportions do not add up to 1, it signifies that there is an additional unknown component.

Table S8: reAdmix using a database of ancient individuals. A maximum of four source populations was allowed for each sample. When proportions do not add up to 1, it signifies that there is an additional unknown component.

