## Supplemental Material for "Diverse genetic origins of medieval steppe nomad conquerors"

**Samples**

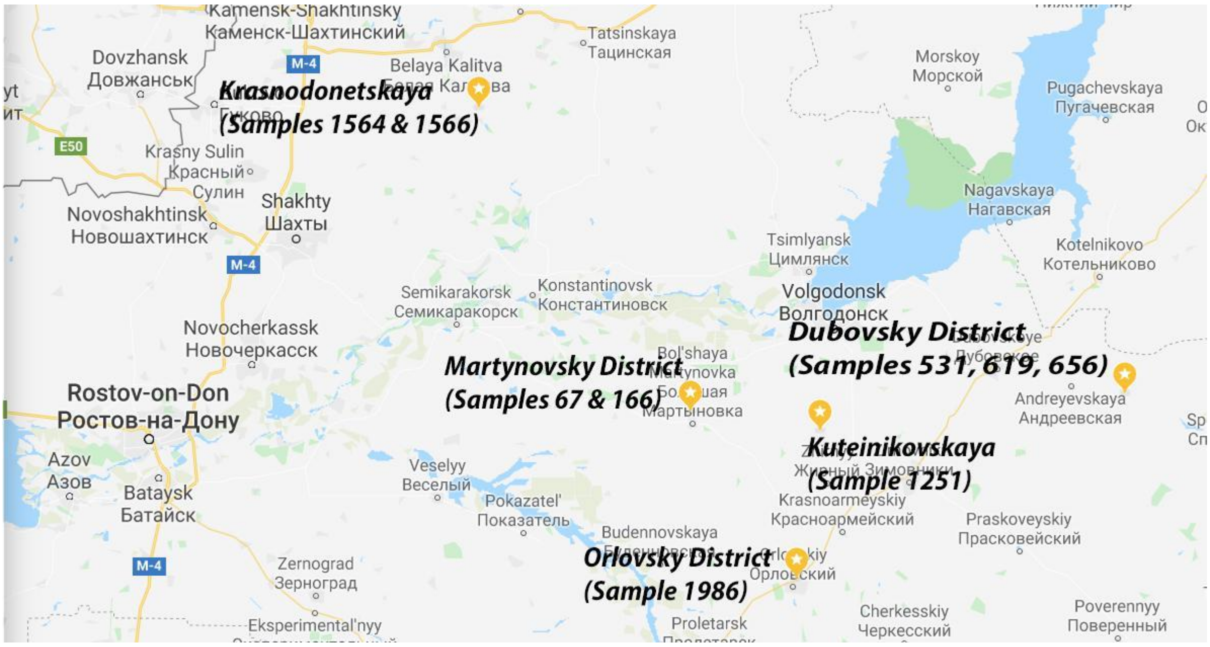


*Figure SF1: Location of the kurgans and samples.*

The nine skeletons were collected from typical Khazar burial kurgans in the southern Russian steppes. Upper-class Khazar burials were square (up to 40 meters) relatively low (presently less than one meter tall) mounds surrounded by ritual trenches (one meter wide and 1-1.5 meter deep). The ritual trenches were originally filled with bones of sacrificed animals and other religious artifacts. The body was placed at the ground level. Sometimes the human body was accompanied by bodies of service animals (horses and camels). Jewelry, weapons, and food for the afterlife (frequently placed in ceramic jars) were also placed inside the mound. Since the mounds were made of earth (without, or almost without, stone foundations) as time passed, the kurgans were flattened, and the shape became nearly circular. Therefore, it is very difficult to determine if the original kurgan was square or if it was a square burial inside the round kurgan. Sometimes, there were multiple graves within the same mound that can be attributed to members of the same clan of a family. Archaeologists can distinguish between primary and secondary burials based on the positions of bodies within the mound. Kurgans frequently occurred in sets, for example, along the river banks.

There are two major types of Khazar burials: “stepped graves” (a rectangular pit with steps/shoulders along the long side) (Figure SF2, A) and “niche grave”, a vertical pit grave with a horizontal niche in the south wall (Figure SF2, B). The second type of grave is the most common for Khazars.

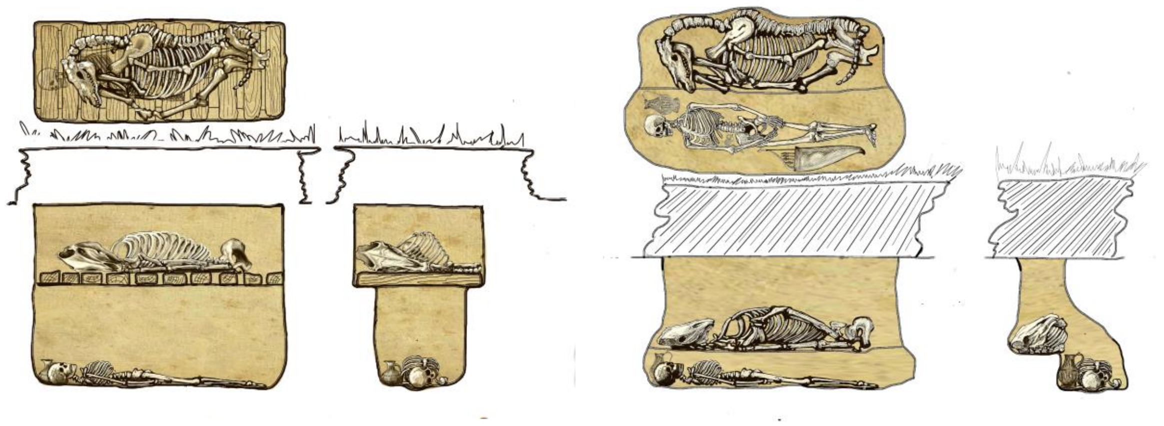

*Figure SF2: Artistic rendering of stepped (A) and niche graves (B).*

Early Khazar burials frequently contained a skeleton of a horse. Later burials contained only inedible parts of a horse (a horse representation made from head, hooves, and skin).

**Sample 1251**

This sample was from The Kuteynikovsky burial mound II that was excavated in 1994 near the village of Kuteinikovskaya Zimovnikovsky district of the Rostov region by the Novocherkassk archaeological expedition.

In the primary burial of Kurgan 2, a male body was uncovered, the age is estimated to be over 40 years. The skull was missing. The grave is in the center of the square surrounded by a trench, filled with horses’ bones and ten horse skulls without lower jaws. The grave is of the pit type with a niche in the southern wall. The grave was robbed; however, some artifacts and their fragments were left behind: a silver belt buckle and silver ornamented tip of a belt, an iron knife, a fragment of an iron object (possibly a saber), and a round pot with a linear-wavy ornament. The buried man apparently suffered from advanced-stage osteochondrosis, with several vertebrae fused, suggesting that he could have been a hunchback. The burial is dated to the late VII$^{th}$ - early VIII$^{th}$ century ("Early Khazar" period), which is supported by the archaic type of the connector of the belt buckle - with a fixed jumper under the tongue. A more advanced flexible connection, with a pin threaded through the hole at the end of the flap buckles, was not developed until the VIII century. Burials with square trenches of this time period are extremely rare. This funerary custom became widespread in the second half of the VIII century.

**Sample 1564**

Nizhnedonsky Kurgans (Nizhnedonsky Chastye Kurgany) are located at the village of Krasnodonetskaya, Belokalitvinsky district of the Rostov region. The mound was excavated in 2001 by the expedition of the Southern Federal University. Sample 1564 was the primary burial in Kurgan 3. It belonged to a male, aged 25 - 35 years. The grave was located in the center of the site surrounded by a square trench. The grave is of the pit type with a niche in the southern wall. It was robbed, leaving several items behind, such as horseshoe-shaped ornamental belt overlay, in the "Crimean-Byzantine" style, and fragments of ivory bow overlays. The burial is from late VIII$^{th}$ to the early IX$^{th}$ century.

**Sample 166**

This family of Kurgans is in the Martynivsky district of the Rostov region; it was excavated in 1982. Sample 166 is the primary burial in Kurgan 37. The remains were of a female, aged 25 - 30 years. The grave is of the pit type with a niche in the southern wall. Fortunately, the grave was not robbed and contained multiple items, including pieces of ironwork and stirrups. The burial is dated between the second half of the VIII$^{th}$ century and early IX$^{th}$ century.

**Sample 1566**

Nizhnedonsky Kurgans (Nizhnedonsky Chastye Kurgany) are located at the village of Krasnodonetskaya, Belokalitvinsky district of the Rostov region. They were excavated in 2001 by the expedition of the Southern Federal University. Sample 1566 is the primary burial from Kurgan 9. The grave belonged to a male, aged 35 - 40 years. The grave was of the stepped type. It was robbed, leaving: several three-lobed arrowheads, a gold earring with a bead, fragments of iron objects, and possible parts of a quiver. The burial is dated between the second half of the VIII century and early IX$^{th}$ century.

**Sample 1986**

Kurgan burial ground Talovy II is in the Orlovsky district of the Rostov region. It was excavated in 2004 by the expedition of the Archaeological Research Bureau, Rostov-on-Don. Sample 1986 was the primary grave of Kurgan 3. It belonged to a male, aged 35 - 45 years. The man was buried along with a horse and a camel. The grave is of the pit type with niches in the southern and western walls. The burial was not robbed, providing multiple artifacts, including arrowheads, ivory bow overlays, and a flail. The main finding is a reliquary made from a carved deer antler, showing combat scenes between a rider and a foot soldier. Considering the item's complexity and level of craftsmanship, it is a rare find worldwide, and the only one found in the Lower Don area. The burial is dated between the second half of the VIII$^{th}$ - and early IX$^{th}$ century.

**Sample 531**

This sample was excavated from Podgornensky IV site, mound 22, burial 1. This kurgan is located near the Podgornensky farm, Dubovsky district of the Rostov region, on the left bank of the Don River (now the left bank of the Tsimlyansk reservoir). In the kurgan, at least five Khazar time burials related to the square trenches culture were found. The trenches were clearly identified in four out of five burials. Kurgan 22 was excavated by the 1988 expedition of Rostov State University [1]. The height of the kurgan was 0.2 m. It was surrounded by the square trench measuring 9 x 9 m. The first burial was made of the stepped style, with a complete horse skeleton placed on the step. Also, the remains of a Caucasoid male 35 - 40 years old at the time of death were found [2]. The site was robbed, however, there were several artifacts remaining. Iron bits were found with S-shaped psalms and arch-shaped iron stirrups with a trapezoid loop of the clutch and a curved footboard. There was also a fragment of a container from the horn (apparently the neck of a water-skull) on the surface of which there are several signs of runic writing [3]. Dating of the burial: VIII$^{th}$ - the first third of the IX$^{th}$ centuries. Information about this burial has not been published in the scientific literature.

**Sample 619**

The sample is from Podgornensky V site, kurgan 5, burial 1. The kurgan is located near the Podgornensky farm, Dubovsky district of the Rostov region, on the left bank of the Don (now the left bank of the Tsimlyansk reservoir). In the burial site, at least two Khazar period burials related to the square trench culture were found. The trench was identified in one of two kurgans. Kurgan 5 was excavated by the expedition of the Rostov State University in 1988 [1]. The height of the kurgan was 0.33 m. A square trench with dimensions of 12.5 x 12.5 m was present, as well as a u-shaped trench with an extension of 6.5 x 9.7 m, which adjoined the main trench from the north. Two burials were found in the kurgan: one of them was in the center of the kurgan marked by a square trench, the other was in the center of the location marked by the u-shaped trench. Burial 1 is of the pit type with a niche in the southern wall. An incomplete skeleton of the horse, the skull and the limbs, was found in the grave. The grave contained one male skeleton (35 - 40 years old), supposedly Mongoloid [2]. The burial was not robbed. Iron bits were found with S-shaped psalms and arch-shaped iron stirrups, as well as a bronze part of a harness. In addition, silver belt fragments were found: an openwork belt buckle and an openwork belt lining of a horseshoe shape. A small golden rectangular plate was also found - probably a detail of a belt leather handbag. Bone facings of onions were also found: middle-end and lateral central ones. On one of them (central side), signs of runic writing are scratched [3]. The dating of the burial: the end of the VII$^{th}$ - early VIII$^{th}$ centuries. These findings have not been fully published.

**Sample 656**

This sample was excavated from the Verbovy log IX site, kurgan 3, burial 1. This site is near the Verbovy Log Farm of Dubovsky district of the Rostov region located on the left bank of the Don (now the left bank of the Tsimlyansk reservoir). Kurgan 3 was excavated by the expedition of the Rostov State University (Rostov-on-Don) in 1989 [1]. The height of the kurgan was 0.24 m. The square below the square was surrounded by a square trench measuring 14.5 x 12 m. Burial 1 is the pit type with a niche in the southern wall. In the grave, there was a male skeleton (30 - 35 years), likely Caucasoid in appearance [4]. The burial has not been robbed.

At the bottom of the grave, there was an incomplete skeleton of the horse, the skull and the limbs. There were iron stirrups, iron buckle, iron arrowheads, iron quiver hook, quiver loops, fragments of iron blade weapons (probably a broadsword with a handle decorated with silver lining), and with ivory lining. An oval-shaped, silver belt buckle was also found [3,5]. In the mouth of the deceased was half of the Byzantine gold coin from the second reign of Emperor Justinian II (705-711 AD). Typical Turkic runes were scratched on the surface of the coin. There were also coin-like discs similar in size to the Byzantine coin [6]). Therefore, the burial is dated to the first third of the VIIIth century, and not before 705. This work has not been fully published.

**Sample 67**

Kurgan burial site Krivolimansky I is located at the Krivoy Liman farm of the Martynovsky district of the Rostov region (Left Bank of the Don River). Excavations were conducted in 1980 by E.I. Savchenko and L.M. Kazakova from Rostov State University [4]. The mound #52 has two burials, and the main burial contained one skeleton of a 35 - 40 years old male, dated to the IX century AD. There was no square trench detected.  The grave is of the pit type with a niche in the southern wall, with the head of the skeleton to the west.  Based on the overall structure and attributes, this burial belongs to the class of burials of mounds with trenches, although the trench itself was not found in this case (maybe the trench was shallow and could not be traced). In general, in Khazar graves, such trench is found in approximately 60% of cases. The buried male was a Mongoloid approximately 35-40 years old. The skull is well preserved. The nose was fractured, and the teeth chipped. The jaw showed signs of inflammation, it was probably broken. There were also damages to the joints of the heads of the humerus and pelvis.

*Table S1: Samples description showing gender, age, date, location, and sequencing statistics (bases covered, average coverage, the fraction of the genome covered by at least one read)*

| Sample | Bases covered | Average coverage | The fraction of genome covered by >=1 read | Bone | Gender | Age at death | Century | Location (district) in the Rostov county | Skull morphology |
|---|---|---|---|---|---|---|---|---|---|
| 67 | 266114863 | 0.1 | 0.09 | Left humerus | M | 35-40 | IX | Martynovsky | Mongoloid |
| 166 | 1.151E+09 | 0.43 | 0.34 | Left femur | F | 25-30 | VIII-IX | Martynovsky | Mongoloid |
| 531 | 56483590 | 0.02 | 0.02 | Right tibia and left ulna | M | 35-40 | VIII-IX | Dubovsky | Caucasoid |
| 619 | 122251586 | 0.05 | 0.04 | Left femur | M | 35-40 | VII-VIII | Dubovsky | Mongoloid |
| 656 | 206775736 | 0.08 | 0.07 | Right tibia | M | 30-35 | VII-VIII | Dubovsky | Caucasoid |
| 1251 | 1.096E+09 | 0.41 | 0.32 | Left humerus | M | 40 | IX | Zimovnikovsky | undefined (skull missing) |
| 1564 | 729454376 | 0.27 | 0.23 | Left tibia | M | 25-35 | VIII-IX | Belokalitvinsky | Caucasoid |
| 1566 | 1.226E+09 | 0.46 | 0.36 | Right humerus | M | 35-40 | VIII-IX | Belokalitvinsky | Mongoloid |

| | | | | | | | | | |
|---|---|---|---|---|---|---|---|---|---|
| 1986 | 1.463E+09 | 0.55 | 0.41 | Right humerus and left tibia | M | 35-45 | VIII-IX | Orlovsky | Mongoloid |

**Contamination analysis**

We have used ANGSD [7] to estimate contamination levels in the analyzed genomes, using the X chromosome that exists in one copy for male samples. ANGSD conducts Fisher's exact test for finding a p-value and uses the jackknife procedure to estimate contamination for each sample. For control, we have also applied *ANGSD* to the single female sample. We present two estimates: Methods of Moments (MoM) and Maximum Likelihood (ML) discarding and not discarding X chromosomal regions with low mappability. ANGSD results show that the level of contamination is acceptably low, not exceeding 1% (see table S2 (Contamination Analysis) in this supplement). Samples 531 and 619 did not have enough coverage for all (531) or some (619) of the analysis.

*Table S2: Contamination. The table lists the results of ANGSD for 9 Khazarian samples. Two estimators are used: Methods of Moments (MoM) and Maximum Likelihood (ML). Also, the analysis can be conducted discarding (right column) and not discarding (middle column) X chromosomal regions with low mappability [7]. Contamination level is computed by 4 methods (Method 1(old/new) and Method 2(old/new), using the code from Rasmussen et al [7]).*

| **Sample** | **Entire X chromosome** | **X:50 000 000-154 900 000** |
|---|---|---|
| 1251<br>Male | We have nSNP sites: 1170, with flanking: 10530<br>Mismatch_rate_for_flanking:0.002687<br>MisMatch_rate_for_snpsite:0.004370<br><br>Method1: old_llh Version: MoM:0.003573<br>SE(MoM):3.467529e-03 ML:0.007886<br>SE(ML):8.481517e-15<br>Method1: new_llh Version: MoM:0.003560<br>SE(MoM):3.480064e-03 ML:0.007844<br>SE(ML):4.033169e-15<br>Method2: old_llh Version: MoM:0.006703<br>SE(MoM):5.918608e-03 ML:0.010064<br>SE(ML):1.186226e-15<br>Method2: new_llh Version: MoM:0.006678<br>SE(MoM):5.941270e-03 ML:0.010009<br>SE(ML):4.270414e-15 | We have nSNP sites: 696, with flanking: 6264<br>Mismatch_rate_for_flanking:0.002518<br>MisMatch_rate_for_snpsite:0.004689<br><br>Method1: old_llh Version: MoM:0.004844<br>SE(MoM):4.636470e-03 ML:0.009248<br>SE(ML):1.829290e-16<br>Method1: new_llh Version: MoM:0.004828<br>SE(MoM):4.651416e-03 ML:0.009205<br>SE(ML):1.143306e-15<br>Method2: old_llh Version: MoM:0.004844<br>SE(MoM):6.962706e-03 ML:0.008395<br>SE(ML):1.509165e-15<br>Method2: new_llh Version: MoM:0.004828<br>SE(MoM):6.987083e-03 ML:0.008351<br>SE(ML):5.487871e-16 |
| 1566<br>Male | We have nSNP sites: 1094, with flanking: 9846<br>Mismatch_rate_for_flanking:0.000593<br>MisMatch_rate_for_snpsite:0.003020 | We have nSNP sites: 678, with flanking: 6102<br>Mismatch_rate_for_flanking:0.000607<br>MisMatch_rate_for_snpsite:0.004861 |

| | | |
|---|---|---|
| | Method1: old_llh Version: MoM:0.006725 SE(MoM):3.201532e-03 ML:0.006970 SE(ML):4.444696e-15<br>Method1: new_llh Version: MoM:0.006719 SE(MoM):3.203922e-03 ML:0.006965 SE(ML):9.749655e-16<br>Method2: old_llh Version: MoM:0.008395 SE(MoM):5.033391e-03 ML:0.008644 SE(ML):3.441055e-16<br>Method2: new_llh Version: MoM:0.008388 SE(MoM):5.036255e-03 ML:0.008640 SE(ML):1.376422e-15 | Method1: old_llh Version: MoM:0.011868 SE(MoM):5.149756e-03 ML:0.011962 SE(ML):3.881710e-15<br>Method1: new_llh Version: MoM:0.011858 SE(MoM):5.153657e-03 ML:0.011954 SE(ML):3.610893e-15<br>Method2: old_llh Version: MoM:0.014575 SE(MoM):8.161987e-03 ML:0.015126 SE(ML):6.950968e-15<br>Method2: new_llh Version: MoM:0.014563 SE(MoM):8.171447e-03 ML:0.015105 SE(ML):2.256808e-16 |
| 1564 Male | We have nSNP sites: 535, with flanking: 4815<br>Mismatch_rate_for_flanking:0.001985<br>MisMatch_rate_for_snpsite:0.005282<br><br>Method1: old_llh Version: MoM:0.007726 SE(MoM):5.725068e-03 ML:0.008415 SE(ML):6.814748e-16<br>Method1: new_llh Version: MoM:0.007703 SE(MoM):5.741185e-03 ML:0.008409 SE(ML):1.122429e-15<br>Method2: old_llh Version: MoM:0.005040 SE(MoM):7.808870e-03 ML:0.004672 SE(ML):1.122429e-15<br>Method2: new_llh Version: MoM:0.005025 SE(MoM):7.829864e-03 ML:0.004665 SE(ML):5.411712e-16 | We have nSNP sites: 341, with flanking: 3069<br>Mismatch_rate_for_flanking:0.001918<br>MisMatch_rate_for_snpsite:0.004178<br><br>Method1: old_llh Version: MoM:0.005550 SE(MoM):6.572008e-03 ML:0.005363 SE(ML):5.597676e-16<br>Method1: new_llh Version: MoM:0.005534 SE(MoM):6.589133e-03 ML:0.005357 SE(ML):0.000000e+00<br>Method2: old_llh Version: MoM:0.002826 SE(MoM):8.340314e-03 ML:0.006315 SE(ML):3.518539e-16<br>Method2: new_llh Version: MoM:0.002818 SE(MoM):8.352485e-03 ML:0.006303 SE(ML):7.996680e-17 |
| 166 Female | We have nSNP sites: 3261, with flanking: 29349<br>Mismatch_rate_for_flanking:0.000878<br>MisMatch_rate_for_snpsite:0.083552<br><br>Method1: old_llh Version: MoM:0.237141 SE(MoM):8.662802e-03 ML:0.236121 SE(ML):5.261349e-13<br>Method1: new_llh Version: MoM:0.236822 SE(MoM):8.674148e-03 ML:0.235835 SE(ML):3.343809e-13<br>Method2: old_llh Version: MoM:0.245480 SE(MoM):1.392879e-02 ML:0.243317 SE(ML):8.335752e-13<br>Method2: new_llh Version: MoM:0.245151 SE(MoM):1.394597e-02 ML:0.243000 SE(ML):1.074456e-12 | We have nSNP sites: 2040, with flanking: 18360<br>Mismatch_rate_for_flanking:0.000767<br>MisMatch_rate_for_snpsite:0.083443<br><br>Method1: old_llh Version: MoM:0.237188 SE(MoM):1.097685e-02 ML:0.237200 SE(ML):1.817301e-13<br>Method1: new_llh Version: MoM:0.236906 SE(MoM):1.098775e-02 ML:0.236946 SE(ML):2.055430e-13<br>Method2: old_llh Version: MoM:0.225384 SE(MoM):1.694188e-02 ML:0.224604 SE(ML):1.955165e-13<br>Method2: new_llh Version: MoM:0.225116 SE(MoM):1.695993e-02 ML:0.224332 SE(ML):1.854900e-13 |
| 1986 Male | We have nSNP sites: 1518, with flanking: 13662<br>Mismatch_rate_for_flanking:0.000844 | We have nSNP sites: 958, with flanking: 8622<br>Mismatch_rate_for_flanking:0.000916 |

| | | |
|---|---|---|
| | MisMatch_rate_for_snpsite:0.002149<br><br>Method1: old_llh Version: MoM:0.003823<br>SE(MoM):2.415519e-03 ML:0.004323<br>SE(ML):4.189042e-15<br>Method1: new_llh Version: MoM:0.003818<br>SE(MoM):2.418314e-03 ML:0.004318<br>SE(ML):6.013303e-15<br>Method2: old_llh Version: MoM:0.006929<br>SE(MoM):4.247989e-03 ML:0.006717<br>SE(ML):2.702608e-16<br>Method2: new_llh Version: MoM:0.006920<br>SE(MoM):4.253614e-03 ML:0.006708<br>SE(ML):3.378260e-16 | MisMatch_rate_for_snpsite:0.001951<br><br>Method1: old_llh Version: MoM:0.002886<br>SE(MoM):2.887446e-03 ML:0.003470<br>SE(ML):4.829795e-16<br>Method1: new_llh Version: MoM:0.002882<br>SE(MoM):2.891255e-03 ML:0.003465<br>SE(ML):7.647176e-16<br>Method2: old_llh Version: MoM:0.006407<br>SE(MoM):5.292941e-03 ML:0.007773<br>SE(ML):3.810172e-15<br>Method2: new_llh Version: MoM:0.006398<br>SE(MoM):5.298761e-03 ML:0.007763<br>SE(ML):6.010412e-15 |
| 531<br>Male | We have nSNP sites: 64, with flanking: 576<br>Not enough SNPs | We have nSNP sites: 34, with flanking: 306<br>Not enough SNPs |
| 619<br>Male | We have nSNP sites: 244, with flanking: 2196<br>Mismatch_rate_for_flanking:0.000252<br>MisMatch_rate_for_snpsite:0.000000<br><br>Method1: old_llh Version: MoM:-0.000675<br>SE(MoM):6.776616e-04 ML:-0.000269<br>SE(ML):4.225260e-18<br>Method1: new_llh Version: MoM:-0.000674<br>SE(MoM):6.778950e-04 ML:-0.000269<br>SE(ML):1.098568e-17<br><br>We have nSNP sites: 252, with flanking: 2268<br>Mismatch_rate_for_flanking:0.000244<br>MisMatch_rate_for_snpsite:0.000000<br><br>Method1: old_llh Version: MoM:-0.000658<br>SE(MoM):6.606005e-04 ML:-0.000261<br>SE(ML):6.011947e-18<br>Method1: new_llh Version: MoM:-0.000657<br>SE(MoM):6.608235e-04 ML:-0.000261<br>SE(ML):6.011947e-18 | We have nSNP sites: 146, with flanking: 1314<br>Not enough SNPs |
| 656<br>Male | We have nSNP sites: 187, with flanking: 1683<br>Mismatch_rate_for_flanking:0.000659<br>MisMatch_rate_for_snpsite:0.002639<br><br>Method1: old_llh Version: MoM:0.005576<br>SE(MoM):7.529508e-03 ML:0.006226<br>SE(ML):4.731695e-17<br>Method1: new_llh Version: MoM:0.005571<br>SE(MoM):7.535962e-03 ML:0.006220 | We have nSNP sites: 119, with flanking: 1071<br>Mismatch_rate_for_flanking:0.000517<br>MisMatch_rate_for_snpsite:0.000000<br><br>Method1: old_llh Version: MoM:-0.001358<br>SE(MoM):1.371018e-03 ML:-0.000570<br>SE(ML):0.000000e+00<br>Method1: new_llh Version: MoM:-0.001357<br>SE(MoM):1.371963e-03 ML:-0.000570 |

| | | |
|---|---|---|
| | SE(ML):1.182924e-16<br>Method2: old_llh Version: MoM:0.012976<br>SE(MoM):1.494426e-02 ML:0.013712<br>SE(ML):2.365847e-17<br>Method2: new_llh Version: MoM:0.012964<br>SE(MoM):1.495688e-02 ML:0.013698<br>SE(ML):8.990220e-16 | SE(ML):0.000000e+00<br>Method2: old_llh Version: MoM:-0.001358<br>SE(MoM):2.787243e-03 ML:-0.001158<br>SE(ML):2.591039e-17<br>Method2: new_llh Version: MoM:-0.001357<br>SE(MoM):2.791154e-03 ML:-0.001158<br>SE(ML):2.355490e-18 |
| 67<br>Male | We have nSNP sites: 217, with flanking: 1953<br>Mismatch_rate_for_flanking:0.000281<br>MisMatch_rate_for_snpsite:0.000000<br><br>Method1: old_llh Version: MoM:-0.000758<br>SE(MoM):7.622029e-04 ML:-0.000296<br>SE(ML):2.310496e-17<br>Method1: new_llh Version: MoM:-0.000758<br>SE(MoM):7.625004e-04 ML:-0.000296<br>SE(ML):2.310496e-17<br>Method2: old_llh Version: MoM:-0.000758<br>SE(MoM):1.564031e-03 ML:-0.000607<br>SE(ML):1.593445e-17<br>Method2: new_llh Version: MoM:-0.000758<br>SE(MoM):1.565284e-03 ML:-0.000607<br>SE(ML):2.071479e-17 | We have nSNP sites: 138, with flanking: 1242<br>Mismatch_rate_for_flanking:0.000442<br>MisMatch_rate_for_snpsite:0.000000<br><br>Method1: old_llh Version: MoM:-0.001151<br>SE(MoM):1.160698e-03 ML:-0.000499<br>SE(ML):1.269026e-17<br>Method1: new_llh Version: MoM:-0.001151<br>SE(MoM):1.161367e-03 ML:-0.000499<br>SE(ML):1.269026e-17<br>Method2: old_llh Version: MoM:-0.001151<br>SE(MoM):2.377263e-03 ML:-0.001022<br>SE(ML):2.030442e-17<br>Method2: new_llh Version: MoM:-0.001151<br>SE(MoM):2.380071e-03 ML:-0.001022<br>SE(ML):2.538052e-18 |

**Mitochondrial analysis**

Analysis of mtDNA was conducted using the *BAM Analysis Kit* (http://www.y-str.org/2014/04/bam-analysis-kit.html). Reference population frequencies were obtained from https://www.familytreedna.com**.** Five of the samples have D4 or C4 haplogroups, that are common in modern Siberian populations, such as Evens and Evenks [8,9]. Three samples had haplogroup H (H1a3, H5b, H13c1), which is predominantly found in Europe. Subclade H1 is common in Russia (13.5%), H5 is the most frequent and diverse in the Western Caucasus, and H13 is present in both Europe and West Asia.

*Table S3: Mitochondrial haplogroups. MT age estimates were taken from Behar et al.[10] and Derenko et al. [11].*

| Sample | Coverage (qualymap) | Haplogroup | MT age | Currently common |
|---|---|---|---|---|
| 67 | 30.6959 | D4e5 | 10,239.2+/-4,619.1 | East Asia, also present in North Asia, Southeast Asia, Amerindian |
| 166 | 62.1109 | C4 | 23-42 kya | Eurasia/Far East |
| 531 | 5.4292 | X2e | 9,748.2 ± 5,150.6 | Turkey, UK |
| 619 | 7.5134 | H1a3 | 3200-6800 | Possibly, Southwest Asia |
| 656 | 30.8611 | C4a1 | 16,029.7 ± 3,638.4 | Eurasia/Far East |
| 1251 | 71.0723 | H5b | 7,894.5 ± 2,399.7 | Southwestern Eurasia, Caucasus |
| 1564 | 86.4395 | H13c1 | H13c: 11,408.1 ± 1,666.2 | Europe |
| 1566 | 31.2939 | D4b1a1a | 2,745.1 ± 3,269.2 | Northeast |
| 1986 | 38.5156 | C4a1c | 4,270.3 ± 2,620.8 | Eurasia/Far East |

**Y-chromosome analysis**

Y haplogroups were determined using two methods: (1) genotyping DNA loci using Yfiler PCR-based system, using the following microsatellite loci: DYS456, DYS389I, DYS390, DYS389II, DYS458, DYS19, DYS385 a/b, DYS393, DYS391, DYS439, DYS635, DYS392, YGATA H4, DYS437, DYS438, DYS448; and (2) using NGS reads and BAM analysis tool. The study of Y-chromosome STR loci in object 166 (left thigh bone) has shown a lack of enzymatic amplification product, which has confirmed that the remains belong to a female.

Using STR loci, we were able to reliably determine Y haplogroups for four samples 619 - Q, 1986 and 1251 - R1a, and 656 - C3. The rest of the samples did not have enough regions covered to reliably determine the haplogroups. Our second approach was based on NGS sequencing and using the BAM analysis tool. Due to low coverage, we were only able to confirm haplogroups for samples 1986 and 1251 (R1a). The rest of the samples did not have enough Y-chromosome reads to reliably identify the haplogroup.

**GPS/reAdmix analysis**

*Table S4: The number of Ancestry Informative Markers based on the sequencing quality cut-offs. An insufficient number of markers retained for DP3 is typical for aDNA studies.*

| **Sample** | **Q20 DP2** | **Q30 DP2** | **Q20 DP3** | **Q30 DP3** |
|---|---|---|---|---|
| 1251 | 7057 | 6715 | 1347 | 1140 |
| 1566 | 7404 | 7115 | 1380 | 1158 |
| 1564 | 3448 | 3274 | 512 | 439 |
| 166 | 6538 | 6289 | 1113 | 927 |
| 1986 | 10049 | 9572 | 2273 | 1886 |
| 656 | 877 | 858 | 57 | 47 |
| 531 | 389 | 385 | 8 | 7 |
| 67 | 1166 | 1152 | 79 | 75 |
| 619 | 1041 | 1036 | 37 | 35 |

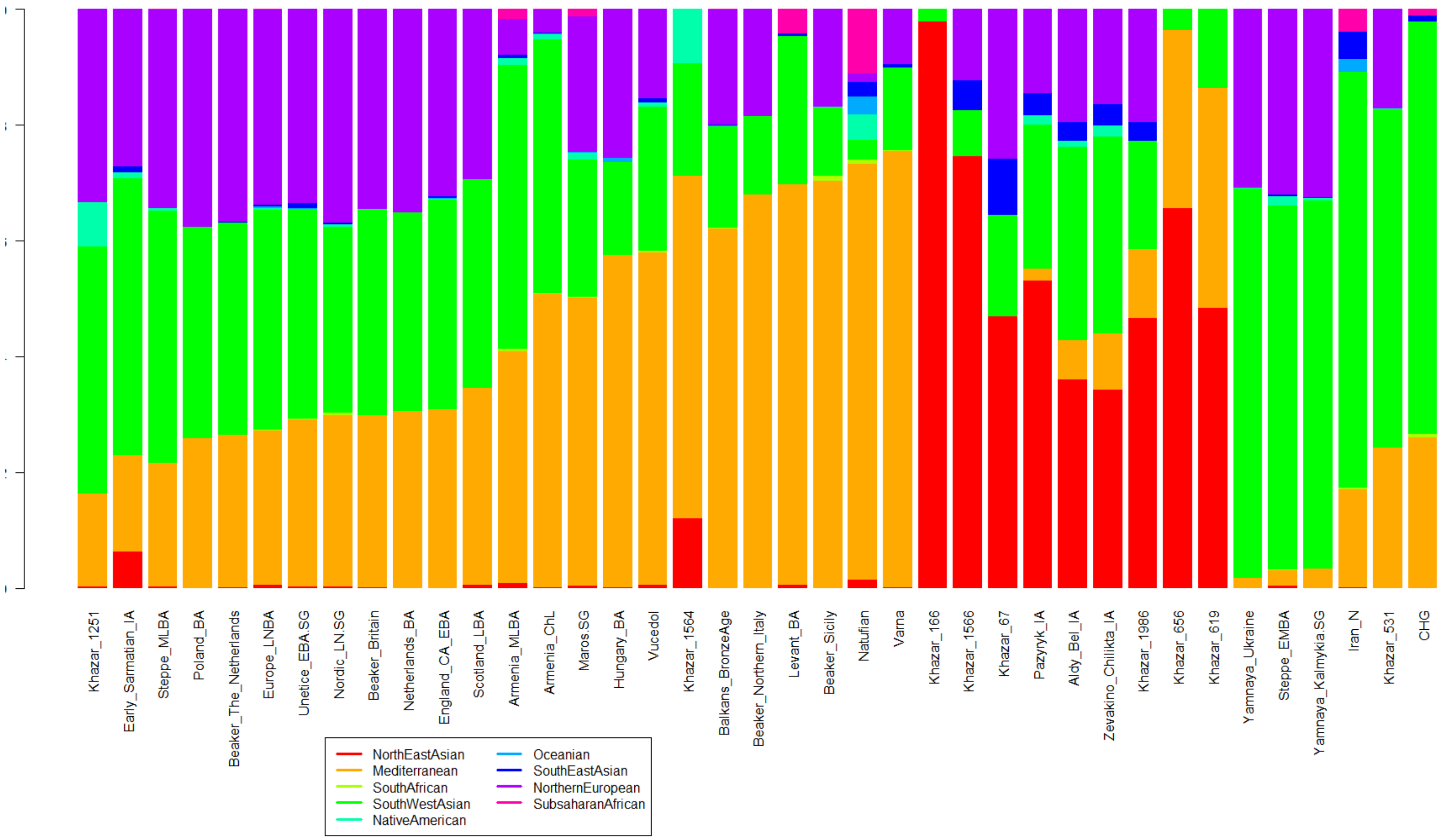


*Figure SF3: Admixture profiles of Khazar samples together with other ancient DNA samples.*

*Table S5: Admixture profiles of Khazar samples for K=9*

| Sample | North-East Asian | Mediterranean | South African | South West Asian | Native American | Oceanian | Southeast Asian | Northern European | Sub-Saharan African |
|---|---|---|---|---|---|---|---|---|---|
| **1251** | 0.004515 | 0.159751 | 1.00E-05 | 0.432111 | 0.074951 | 1.00E-05 | 1.00E-05 | 0.328632 | 1.00E-05 |
| **1564** | 0.121944 | 0.590689 | 1.00E-05 | 0.19407 | 0.093246 | 1.00E-05 | 1.00E-05 | 1.00E-05 | 1.00E-05 |
| **1566** | 0.745746 | 1.10E-05 | 1.00E-05 | 0.078897 | 1.00E-05 | 1.00E-05 | 0.052417 | 0.122889 | 1.00E-05 |
| **166** | 0.976993 | 1.00E-05 | 1.00E-05 | 0.022937 | 1.00E-05 | 1.00E-05 | 1.00E-05 | 1.00E-05 | 1.00E-05 |
| **1986** | 0.466741 | 0.121944 | 1.00E-05 | 0.185399 | 1.00E-05 | 1.00E-05 | 0.033167 | 0.192709 | 1.00E-05 |
| **531** | 1.00E-05 | 0.228597 | 1.00E-05 | 0.582869 | 1.00E-05 | 1.00E-05 | 1.00E-05 | 0.188474 | 1.00E-05 |
| **619** | 0.480255 | 0.38177 | 1.00E-05 | 0.137916 | 1.00E-05 | 1.00E-05 | 1.00E-05 | 1.00E-05 | 1.00E-05 |
| **656** | 0.656524 | 0.30707 | 1.00E-05 | 0.036346 | 1.00E-05 | 1.00E-05 | 1.00E-05 | 1.00E-05 | 1.00E-05 |
| **67** | 0.465869 | 1.00E-05 | 1.00E-05 | 0.181063 | 1.00E-05 | 1.00E-05 | 0.097103 | 0.255915 | 1.00E-05 |

Next, we conducted the GPS analysis, using the samples from the first millennium BC as a reference. Most of the sequenced Khazar samples show similarity to bronze and iron age steppe samples. None of the matches was perfect (distance to the reference population >0.05), and, therefore, we cannot consider any of these samples to be direct descendants of a single currently sequenced ancient cultures - they rather appear to be a mix of multiple ethnic components. To estimate proportions of admixture in the nine samples, we used the reAdmix algorithm [12] and the same reference database of modern and ancient individuals. Using the database of modern individuals, individuals can be represented as shown below. Samples 166, 1686, 67, and 1566 show more Asian influence, samples 1564, 531 and 1251 are of mostly European ancestry with Asian admixture, while samples 619 and 656 appear to be a mosaic of Asian and European origins.

According to craniofacial analysis, Pazyryk crania were similar to Central Asia groups, especially the Kazakh and Uzbek samples [13-16]. The Pazyryk mummies were buried in kurgans similar to Scythian burials found in Ukraine. It is accepted that the nomadic people from the Altai Mountains spread over the Eurasian Steppe in the 1st millennium BC [17].

*Table S6: Summary of GPS results, indicating the nearest reference population and Euclidean distance between the sample and reference admixture vector. A small distance indicates that the reference population is a good match for the sample vector. Large distance may indicate either the absence of a suitable reference population or mixed origin of the sample.*

| SAMPLE | NEAREST MODERN | DISTANCE | NEAREST ANCIENT | DISTANCE |
|---|---|---|---|---|
| **1251** | Tajik | 0.18 | Steppe MLBA | 0.06 |
| **1564** | Lebanese | 0.16 | Levant BA | 0.19 |
| **1566** | Yakut | 0.04 | Pazyryk IA (Altai) | 0.27 |
| **166** | Evenk | 0.09 | Pazyryk IA (Altai) | 0.52 |

| | | | | |
|---|---|---|---|---|
| **1986** | Shor | 0.10 | Pazyryk IA (Altai) | 0.14 |
| **531** | Ishkasim | 0.19 | Early Sarmatian IA | 0.17 |
| **619** | Turkmen | 0.26 | Pazyryk IA (Altai) | 0.40 |
| **656** | Kazakh | 0.29 | Pazyryk IA (Altai) | 0.41 |
| **67** | Khanty | 0.12 | Pazyryk IA (Altai) | 0.16 |

*Table S7: reAdmix using a database of modern individuals. A maximum of four source populations was allowed for each sample. When proportions do not add up to 1, it signifies that there is an additional unknown component.*

| **Sample** | **Populations** | | | | **Proportions** | | | |
|---|---|---|---|---|---|---|---|---|
| **1251** | Turkmen | Abkhazian | Belarusian | Yizu | 0.494 | 0.142 | 0.238 | 0.127 |
| **1564** | Druze | | | | 0.649 | | | |
| **1566** | Yakut | | | | 1 | | | |
| **166** | Yakut | Even Sakha | | | 0.637 | 0.339 | | |
| **1986** | Yakut | Saami | Abkhaz | | 0.368 | 0.428 | 0.167 | |
| **531** | Yaghnobi (Tajikistan) | Kets | Kurmi | Selkup | 0.506 | 0.053 | 0.202 | 0.239 |
| **619** | Egypt | Yakut | Azeri | Yizu | 0.315 | 0.183 | 0.206 | 0.297 |
| **656** | Mongolian | Even Sakha | Egypt | Yizu | 0.369 | 0.252 | 0.193 | 0.187 |
| **67** | Yakut | | | | 0.623 | | | |

*Table S8: reAdmix using a database of ancient individuals. A maximum of four source populations was allowed for each sample. When proportions do not add up to 1, it signifies that there is an additional unknown component.*

| **Sample** | **Populations** | | | | **Proportions** | | | |
|---|---|---|---|---|---|---|---|---|
| **1251** | Steppe Eneolithic | Anatolia Neolithic | SE Iberia CA | Zevakino Chilikta IA | 0.624 | 0.149 | 0.124 | 0.103 |
| **1564** | Peloponnese Neolithic | | | | 0.595 | | | |
| **1566** | Pazyryk IA | | | | 1.000 | | | |
| **166** | Pazyryk IA | | | | 1.000 | | | |
| **1986** | Pazyryk IA | | | | 0.832 | | | |
| **531** | Beaker Central Europe | Armenia MLBA | Yamnaya Ukraine | Maros.SG | 0.388 | 0.217 | 0.258 | 0.138 |
| **619** | Beaker Central Europe | Pazyryk IA | Anatolia Neolithic | Peloponnese Neolithic | 0.489 | 0.237 | 0.175 | 0.099 |
| **656** | Beaker Central Europe | Armenia MLBA | Pazyryk IA | | 0.443 | 0.299 | 0.193 | |
| **67** | Pazyryk IA | | | | 0.875 | | | |

.

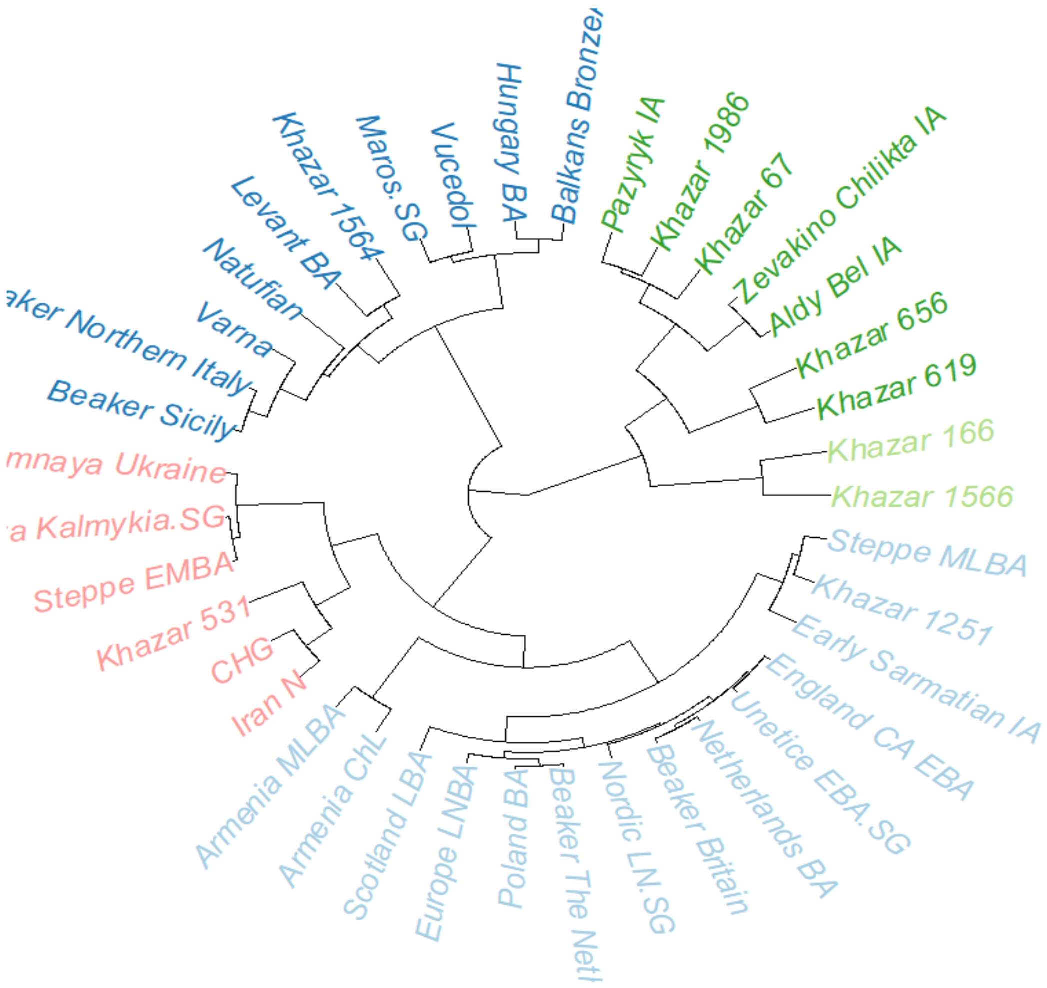


*Figure SF4: Clustering of ancient samples. Six out of nine Khazar samples cluster together (green cluster)*

**TreeMix F3 outgroup analysis**

We have calculated F3 outgroup statistics for the 9 Khazarian samples. Aeta was used as an outgroup. Aeta is an indigenous negrito group living in scattered, isolated mountainous parts of the island of Luzon, the Philippines. According to F3 statistics, two samples (#166 and #1566) were purely Siberian, and the rest of mixed Eastern European origin. However, as suggested by Benjamin Peter [18], the number of pairwise differences may be a better metric for population divergence than the outgroup F3.
